# Structured Chemical Reaction Modeling with Multitask Graph Neural Networks

**DOI:** 10.1101/2025.10.13.682169

**Authors:** Maryam Astero, Anchen Li, Elena Casiraghi, Juho Rousu

## Abstract

Modeling chemical reactions requires connecting fine-grained atom–bond edits with broader semantic categories. Yet, most machine learning approaches model these aspects in isolation: atom mapping, reaction center identification, and reaction classification are treated as separate problems. This separation limits accuracy, interpretability, and generalization. In this work, we argue that *multitask learning* provides a natural and powerful framework for reaction modeling. By jointly predicting mappings, centers, and classes within a single graph neural network, models can leverage structural dependencies between tasks. We proposed MARCC (Mapping-Assisted Reaction Center and Classification), a multitask architecture that achieves state-of-the-art performance on the USPTO-50K benchmark. MARCC demonstrates that multitask supervision not only improves accuracy across all tasks but also provides a structured representation of reaction mechanisms.

## 1 Introduction

Chemical reactions are multi-scale phenomena. At the *local level*, atoms rearrange bonds; at the *intermediate level*, reactive centers define where changes occur; and at the *global level*, transformations fall into semantic classes (e.g., substitution, elimination). Capturing all three perspectives is critical for reaction understanding, template discovery, and mechanistic inference.

Most existing approaches, however, treat these tasks separately. Atom mapping algorithms rely on graph matching or specialized attention modules [1, 2]. Reaction center prediction has been cast as rule-based extraction or subgraph labeling [3, 4]. Reaction classification typically uses fingerprints or graph embeddings [5, 6]. While each area has advanced, the lack of integration overlooks important dependencies: atom mappings, scaffold center prediction, and both local edits and mappings constrain classification.

We argue that multitask learning provides a natural and effective framework for reaction modeling. By jointly predicting mappings, centers, and classes, a multitask approach can exploit structural dependencies between tasks and improve generalization. Beyond performance, it also enhances interpretability by producing a unified structural narrative of the reaction.

First, atom mapping, center prediction, and classification are inherently interdependent. Knowing which atoms align across reactants and products helps pinpoint the true reaction center, which in turn constrains the possible reaction class. Conversely, global reaction type guides which edits are chemically plausible. Ignoring these couplings forces models to rediscover the same signals separately, often inconsistently. Second, multitask supervision acts as a form of regularization. By learning multiple objectives simultaneously, models avoid overfitting to one narrowly defined task. For example, a classifier trained only on reaction classes may exploit dataset biases, whereas coupling it with atom- and bond-level predictions discourages shortcut learning. Finally, multitask outputs support interpretability. By producing mappings, centers, and classes together, the model yields evidence chains: why a bond edit was chosen, how it relates to atom correspondence, and which global class explains the transformation. This structured output aligns naturally with how chemists reason about reactions.

In the following, we introduce MARCC (Mapping-Assisted Reaction Center and Classification), a multitask graph neural framework that operationalizes these ideas by jointly predicting atom mappings, reaction centers, and reaction classes within a unified architecture.

## 2 The MARCC Framework

To study the value of multitask learning in reaction modeling, we introduce MARCC (Mapping-Assisted Reaction Center and Classification), a graph neural network that jointly predicts atom mappings, reaction centers, and reaction classes. Instead of treating these tasks separately, MARCC integrates them into a unified architecture where each task supports the others (Fig. 1).

**Figure 1:**
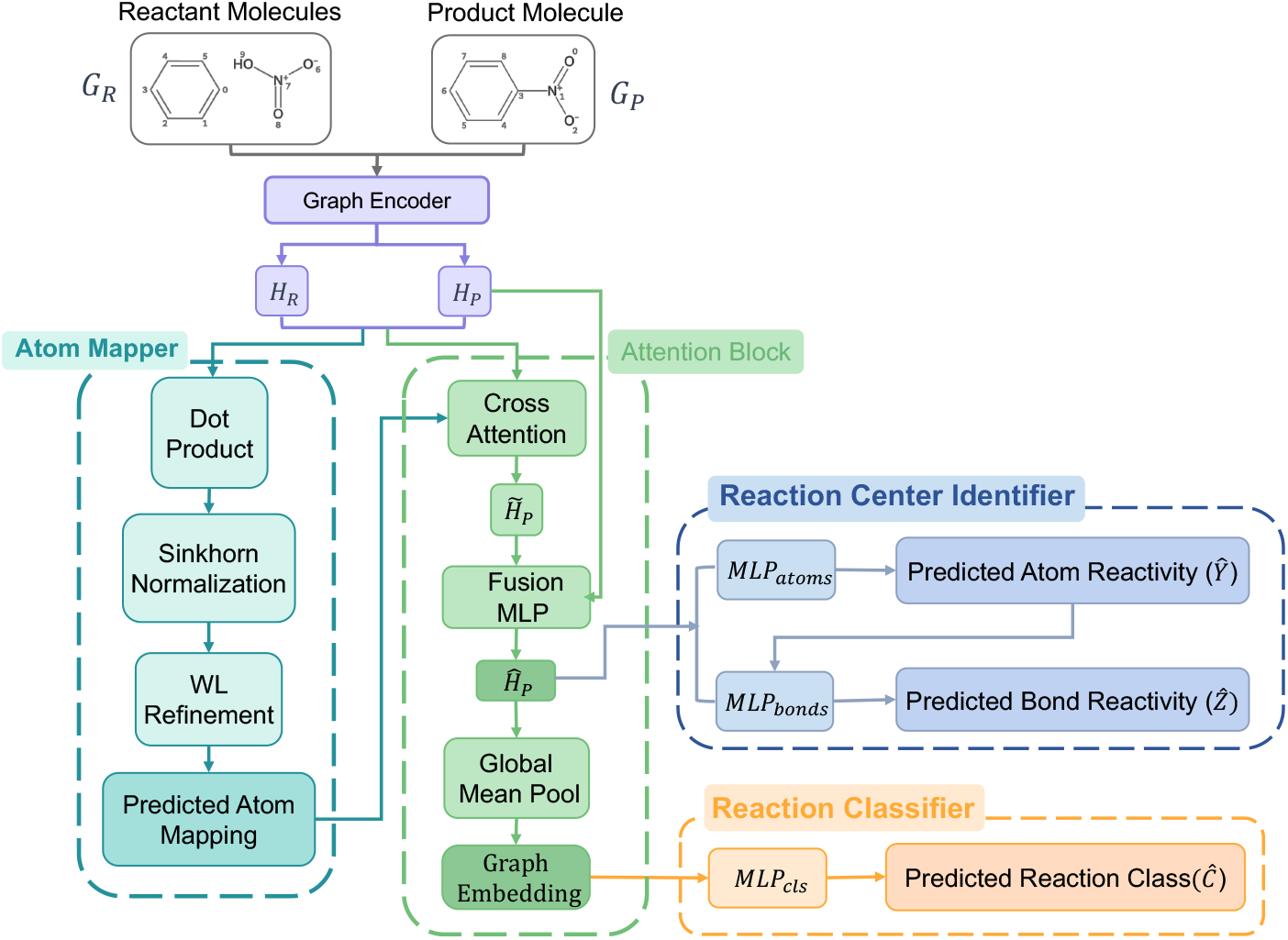
Overview of the MARCC framework. Reactant (*G*_*R*_) and product (*G*_*P*_ ) graphs are encoded with a shared GINE encoder, producing embeddings *H*_*R*_ and *H*_*P*_ . An **Atom Mapper** estimates soft atom correspondences via dot-product similarity and Sinkhorn normalization, refined with Weisfeiler–Lehman (WL) symmetry checks. These alignments guide a cross-attention layer that enriches product embeddings 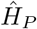, which are then used by task-specific heads for **Reaction Center Identification** and **Reaction Classification**.

### 2.1 Design Principles

MARCC is based on three key ideas:

- **Shared representation:** Reactant and product molecules are encoded by a common GNN, ensuring that all tasks operate over consistent embeddings.
- **Cross-graph alignment:** A differentiable atom-mapping module aligns product atoms to their most likely reactant counterparts. These correspondences are used to guide attention, allowing local edits to be interpreted in the context of structural changes.
- **Task coupling:** Atom mapping provides alignment for reaction center prediction, and both mapping and centers constrain the global reaction class. All tasks share parameters and are optimized jointly with adaptive weighting.

### 2.2 Architecture Overview

MARCC consists of three stages. First, a GNN encoder processes reactant and product graphs into atom- and bond-level embeddings. Second, an atom mapping head estimates soft correspondences between atoms, using Sinkhorn normalization to enforce one-to-one alignment. These alignments inform a cross-attention module that enriches product embeddings with mapped reactant context. Finally, task-specific heads make predictions: binary classifiers label reactive atoms and bonds, while pooled embeddings predict the reaction class.

## 3 Experiments

### 3.1 Experimental Setup

We evaluate MARCC on the USPTO-50K dataset [4, 7, 8], a benchmark with 50k atom-mapped reactions in 10 classes. Canonicalization and remapping [4, 9] ensure index invariance. The dataset is split 8:1:1 into training, validation, and test sets.

Molecular graphs are built from SMILES using RDKit [10], with atom (atomic number, charge, chirality, hybridization, aromaticity, etc.) and bond (bond type, conjugation, stereochemistry) features encoded categorically.

We evaluate three tasks: (i) reaction center prediction (Top-*n* Edit Accuracy), (ii) reaction classification (Top-1 Accuracy), and (iii) atom mapping (symmetry-aware accuracy [11]).

Hyperparameters were optimized with Optuna [12, 13], balancing objectives with uncertainty-based weighting [14]. Training used AdamW (lr=10^*−*4^, wd=5×10^*−*5^), batch size 32, early stopping, and a single NVIDIA V100 GPU.

### 3.2 Impact of Multitask Learning

Table 1 shows the effect of single-task, dual-task, and full multitask configurations.

**Table 1:** Effect of multitask supervision on MARCC. Note: “–” indicates that the corresponding metric was not applicable for the given configuration.

| Configuration | Reaction Center | Classification | Mapping |
| --- | --- | --- | --- |
|  | Edit Acc. (%) | Class Acc. (%) | Map Acc. (%) |
| Classification only | – | 73.3 | – |
| Reactivity only | 87.7 | – | – |
| Mapping only | – | – | 99.0 |
| Classification + React. | 88.1 | 96.3 | – |
| Mapping + Reactivity | 93.0 | – | 90.5 |
| Mapping + Classification | – | 91.8 | 97.6 |
| **Full Multitask** | 99.1 | 97.2 | 98.2 |

Single-task training achieves narrow competence, while dual-task supervision provides partial transfer. The full multitask setup consistently delivers the strongest results, confirming that classification provides global context, reactivity sharpens local edits, and mapping regularizes alignment.

### 3.3 Comparison with State-of-the-Art

We next compare MARCC with leading methods for reaction understanding and atom mapping.

#### Reaction Center and Classification

Table 2 compares MARCC against retrosynthesis baselines (GraphRetro [4], RetroXpert [15], RCSearcher [16]) and classifiers (MolBERT [17], RXGL [6]). We also report a *products-only* variant of MARCC that omits reactants and disables alignment for reaction center prediction, whereas classification always uses both reactants and products.

**Table 2:** Comparison with prior methods on USPTO-50K. Note: “–” indicates that the corresponding metric was not reported in the original study or is not applicable to that method.

| Model | Reaction Center | Classification |
| --- | --- | --- |
|  | Edit Acc. Top-1 (%) | Class Acc. (%) |
| GraphRetro | 70.8 | – |
| RetroXpert | 50.4 | – |
| RCSearcher | 69.3 | – |
| MolBERT | – | 70.3 |
| RXGL | – | 93.2 |
| MARCC (Products only) | 67.9 | 97.6 |
| **MARCC (Full)** | 99.1 | 97.2 |

Results show that MARCC (Full) far outperforms retrosynthesis-focused baselines, achieving near-perfect edit localization and strong classification. The products-only variant, which excludes reactants for center prediction, performs comparably to other product-only baselines. These results suggest that while classification can be solved reliably with both reactants and products, uncovering the true reaction center demands alignment signals, underscoring the benefit of multitask training.

#### Atom Mapping

We further benchmark MARCC’s auxiliary mapping head against dedicated mapping models: RXNMapper [2], GraphormerMapper [18], and SAMMNet [11].

**Table 3:** Atom mapping accuracy on USPTO-50K.

| Model | Accuracy (%) |
| --- | --- |
| RXNMapper | 98.8 |
| SAMMNet | 97.4 |
| GraphormerMapper | 94.5 |
| **MARCC (Full)** | 98.2 |

MARCC achieves 98.2% mapping accuracy, on par with specialized systems, while also excelling in center prediction and classification. This demonstrates the strength of multitask learning, where all three tasks are interdependent and jointly improve one another.

## 4 Conclusion

We demonstrated that multitask learning provides a principled framework for reaction modeling by jointly integrating atom mapping, reaction-center prediction, and reaction classification. Rather than treating these as isolated objectives, MARCC shows that learning them together yields higher accuracy and stronger interpretability. The three tasks reinforce one another: mapping provides alignment, center prediction captures local edits, and classification supplies global semantic context. This synergy not only advances predictive performance but also offers structured, chemically meaningful explanations. Our results highlight multitask learning as a unifying approach to chemical reaction understanding, with broad potential in biochemical and molecular applications.

